# Observation of topological action potentials in engineered tissues

**DOI:** 10.1101/2022.03.16.484369

**Authors:** Hillel Ori, Marc Duque Ramirez, Rebecca Frank Hayward, He Tian, Gloria Ortiz, Adam E. Cohen

**Affiliations:** Department of Chemistry and Chemical Biology, Harvard University, Cambridge, MA 02138; School of Engineering and Applied Science, Harvard University, Cambridge, MA 02138; Department of Chemistry, University of California, Berkeley, CA 94720; Department of Physics, Harvard University, Cambridge, MA 02138

## Abstract

Due to the nonlinear current-voltage relations of ion channels, an interface between two tissues can have very different bioelectrical properties compared to either tissue on its own. Here we show experimentally that gap junction-coupled interfaces between non-excitable tissues can be electrically excitable. This topologically protected excitability occurs over a far larger range of ion channel expression levels than does excitability in the bulk. Topological excitations at tissue interfaces can cause local elevations in calcium concentration, possibly providing a bioelectrical mechanism for interface sensing. As in condensed matter physics, topological excitations in electrophysiology constitute a distinct class of phenomena which may show exotic and novel properties.

## Introduction

Patterns of membrane potential are thought to play a critical role in many biological patterning processes.^1–8^ A natural question is to ask what happens to such patterns at interfaces between tissues whose cells have different resting potentials. When cells are gap junction-coupled across the interface, the voltage along a line transverse to the interface must go through all intermediate values between the two resting potentials. These intermediate voltages may include regions where the membrane potential would be unstable in the bulk due to non-linearity of ion channel current-voltage relations. This continuity requirement could lead to novel excitations localized at tissue interfaces. If two non-excitable tissues are brought into contact, can the interface become excitable?

Interface-localized topological excitations are an emerging area of basic research and applied interest.^9^ Topological excitations have been studied in the context of electronic,^10^ photonic,^11^ mechanical,^12–14^ and hydrodynamic^15^ systems. While these excitations have been widely studied in near-equilibrium systems, biological dynamics provide an opportunity to study topological excitations in far-from-equilibrium nonlinear systems where new types of features may emerge. Recent theoretical work has analyzed the types of topological excitations that might occur in nonequilibrium soft-matter systems,^16–18^ but to our knowledge experimental tests have been lacking, and the possibility of such excitations in bioelectrical systems has not previously been contemplated.

Historically, spatial structures in bioelectrical signaling have been difficult to study because patch clamp measurements probed the voltage at only a single point in space. Recent advances in voltage imaging^19,20^ opened the door to studying the spatial structures of bioelectrical excitations. By combining patterned optogenetic stimulation and high-speed voltage imaging, one can probe the excitability of a complex tissue as a function of space and time. This approach has been used to identify new types of bioelectrical excitations in engineered tissues,^21^ and also to map bioelectrical signals throughout embryonic development.^1,19^ Engineered and patterned cells have been a powerful tool for dissecting intercellular molecular signaling cascades,^22,23^ but despite some theoretical work,^24,25^ this approach has not previously been applied experimentally to bioelectrical signaling.

Here we map the excitability at interfaces between non-excitable engineered tissues and observe interface-localized action potentials. Detailed numerical simulations and a simple analytical model capture the key features of these excitations. Our simulations predict that interface-localized action potentials are exceptionally robust to variations in ion channel levels compared to ordinary action potentials. These findings suggest new mechanisms of bioelectrical signaling that may arise in native tissues and which might be used in engineering synthetic bioelectrical circuits with applications in sensing, tissue engineering, and unconventional computation.

## Results

Human Embryonic Kidney (HEK293) cells become electrically excitable when genetically engineered to express an inward-rectifier potassium channel (e.g. K_ir_2.1 or K_ir_2.3) and a voltage-gated sodium channel (e.g. Na_V_1.3,^29^ Na_V_1.5^30^, or Na_V_1.7^31^). When grown in a confluent monolayer, endogenous gap junctions introduce nearest-neighbor coupling, which can then support propagating electrical waves. If the cells further express a blue light-activated cation channel such as CheRiff^32^ and either express a far-red voltage indicator protein^20^ or are labeled with a red-shifted voltage-sensitive dye^33^, then one can optically induce action potentials and simultaneously map their propagation through the engineered tissue.^21,30,34^

First, we explored whether the sodium and potassium channels needed to be in the same cells. We made separate pools of HEK cells that expressed CheRiff and either Na_V_1.5 alone or K_ir_2.1 alone (Fig. 1A). We mixed the cell populations and mapped the voltage in confluent monolayer cultures using the far-red voltage-sensitive dye BeRST1 (Methods).^33^ Localized optogenetic stimulation evoked action potentials which propagated radially outward (Fig. 1B, Movie 1). In cultures expressing either Na_V_1.5 or K_ir_2.1 alone, optical activation of CheRiff did not lead to action potentials or wave propagation (Fig. S1). These results show that a mixture of non-excitable cells can become excitable by sharing currents through gap junctions.

**Figure 1.**
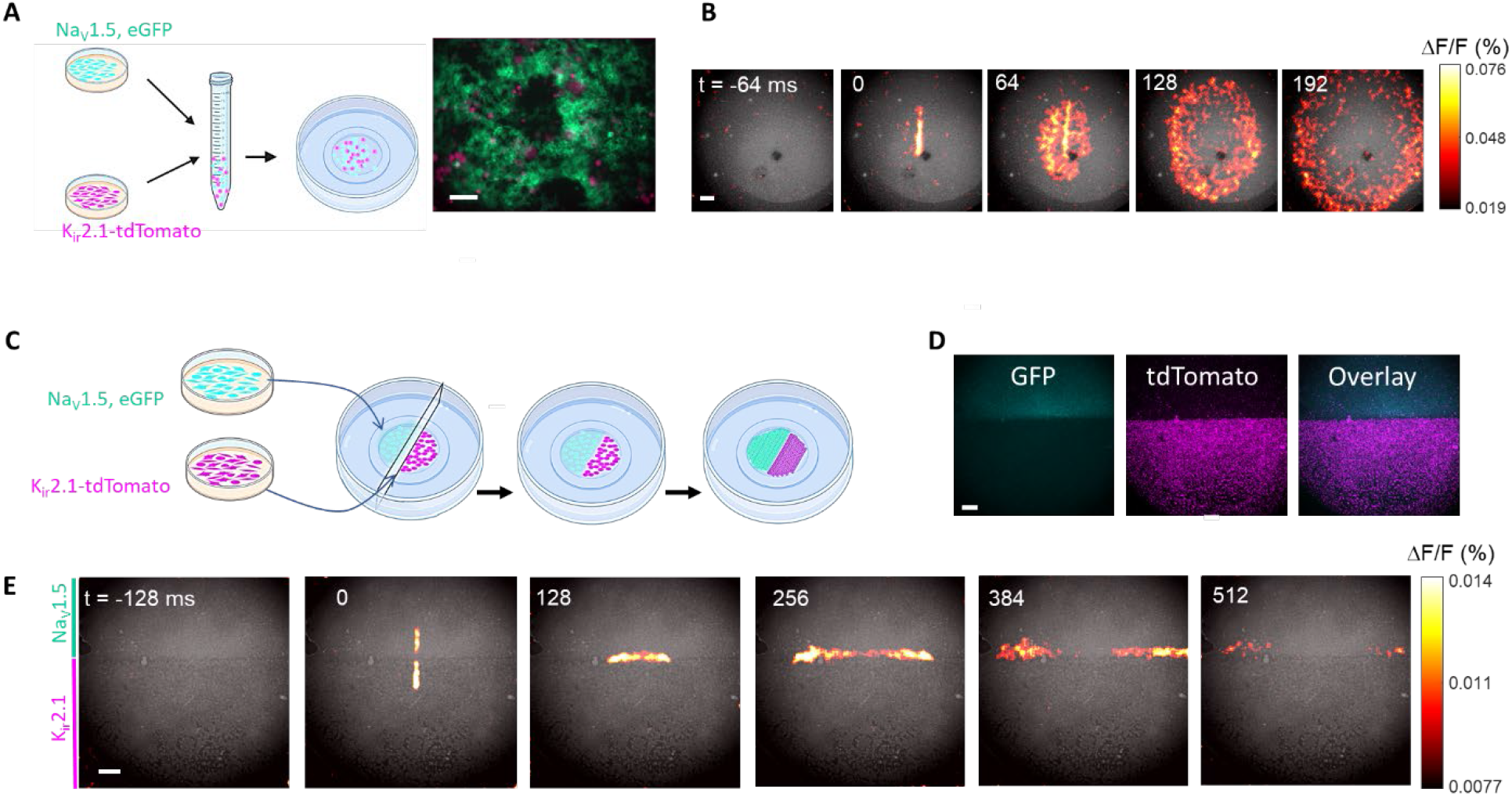
Topological action potentials arise at interfaces of non-excitable tissues. A,B) A mixture of non-excitable cells can be excitable. A) Left: sample preparation. HEK cell lines were engineered to express either K_ir_2.1 (an inward-rectifier K^+^ channel) or Na_V_1.5 (a voltage-gated Na^+^ channel). All cells also expressed a light-gated ion channel, CheRiff. The two populations were mixed, plated, and grown to form a gap junction-coupled monolayer. Right: Distribution of Na_V_1.5-expressing cells (tagged with a GFP marker) and K_ir_2.1-expressing cells (tagged with a tdTomato marker). Expression is mutually exclusive. B) Montage of frames from voltage imaging movie (Movie 1). The monolayer was stimulated by a short pulse of blue light (vertical bar at *t* = 0). Changes in voltage were then monitored via the fluorescence of the far-red voltage-sensitive dye BeRST1 (baseline signal in gray, *ΔF*/*F* in color). Optogenetic stimulation evoked outward propagating action potentials. C-E) Topological action potential at interfaces. C) Sample preparation. HEK cells expressing K_ir_2.1 or Na_V_1.5 were plated on opposite sides of a thin plastic divider. After removal of the divider, the cells migrated to fill the gap and formed a gap junction-coupled interface. D) Interface between populations of Na_V_1.5-expressing cells (tagged with GFP) and K_ir_2.1-expressing cells (tagged with tdTomato). C) Montage of frames from a voltage-imaging movie (Movie 2). Optogenetic stimulation spanning the interface at *t* = 0 evoked a topological action potential that propagated outward solely along the interface (baseline signal in gray, *ΔF*/*F* in color). Scalebars: A) 0.1 mm; B,D,E) 1 mm. See also Figs. S1, S2.

Next, we used a thin plastic divider to physically separate the Na_V_1.5- and the K_ir_2.1-expressing cells on opposite halves of the culture dish (Fig. 1C, Methods). Fluorescent tags marked the two populations (Fig. 1D). After the cells had settled, we gently removed the divider and let the cells migrate to fill the gap (Methods). This approach led to a clean interface with the two cell populations on opposite sides.

Optogenetic stimulation of either population alone did not evoke action potentials. Remarkably, optogenetic stimulation with a stripe that spanned the interface induced electrical excitations which propagated along the interfacial line without entering the bulk on either side (Fig. 1E, Movie 2). The spikes had a width of 0.76 ± 0.15 mm transverse to the interface, a length of 2.21 ± 0.62 mm along the interface, and propagated at 12.9 ± 3.5 mm/s (mean ± s.d., n = 5 samples, Fig. 2A). Thus, topological action potentials can arise between non-excitable tissues. For control experiments, we made interfaces in which one cell population expressed either Na_V_1.5 or K_ir_2.1, and the adjoining population expressed neither channel. These interfaces were not excitable, confirming the necessity of both ion channels on opposite sides of the interface (Fig. S2A,B). Furthermore, in dishes where the Na_V_1.5 and K_ir_2.1-expressing cells had not yet fully closed the gap, topological action potentials propagated only in regions where the two cell types had come into contact (Fig. S2C).

**Figure 2.**
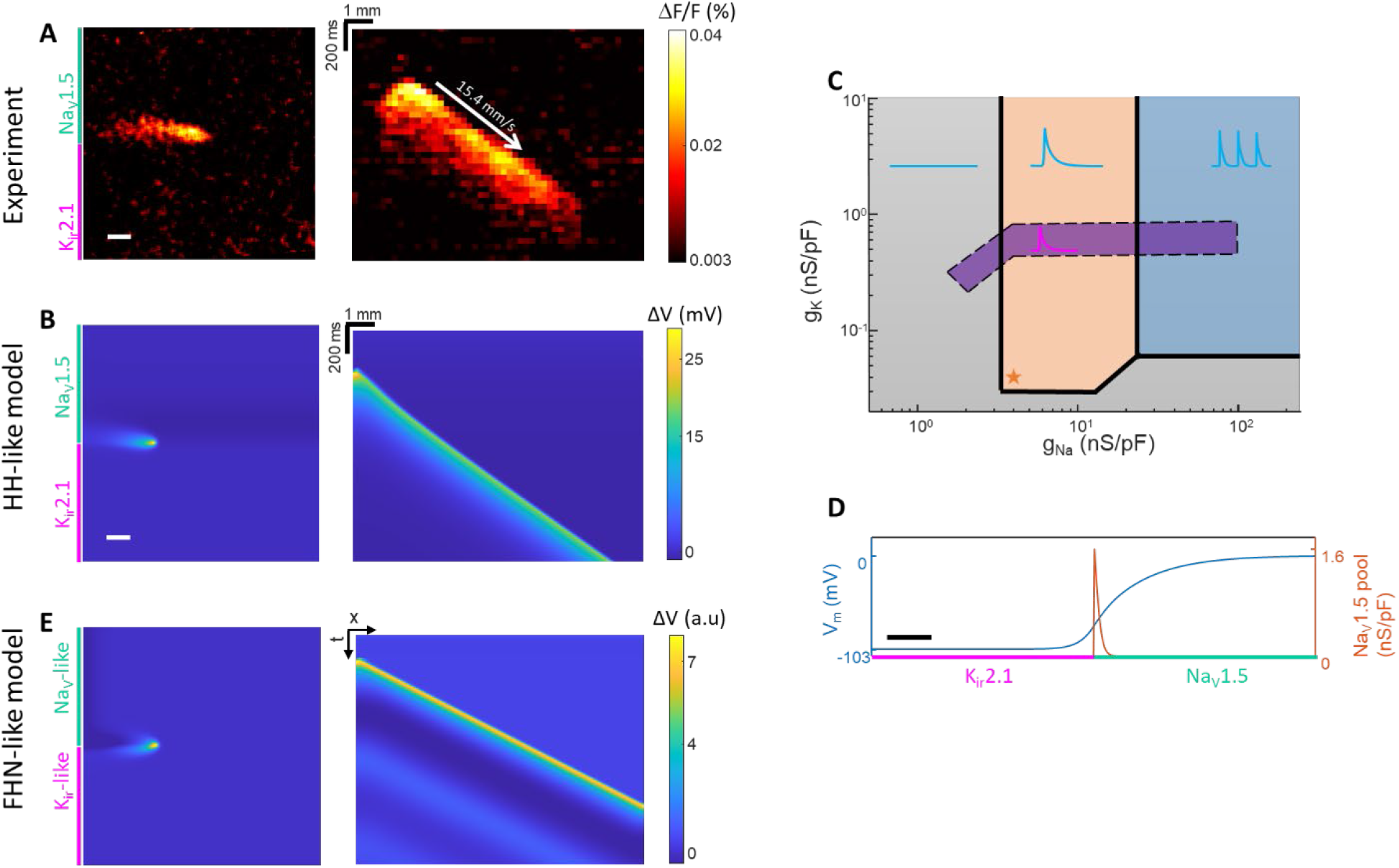
Quantification and numerical simulations of topological action potentials. A) Left: spatial profile of a topological action potential. Right: kymograph constructed by averaging transverse to the interface, showing topological action potential propagation. B) Numerical simulations of an interface between a K_ir_2.1-expressing cell population (g_K_ = 0.04 nS/pF, E_K_ = -107 mV) and a Na_V_1.5-expressing cell population (g_Na_ = 3.9 nS/pF, E_Na_ = 75 mV). Left: Image of a topological action potential propagating along a tissue interface. Color scale corresponds to deviation from local resting potential. Right: kymograph showing topological action potential propagation (Movie 3). C) Calculated phase diagram of excitability of K_ir_2.1-Na_V_1.5 interface. The parameter space is divided into non-excitable, excitable, and spontaneously active phases. The purple region indicates the excitable phase of a homogenous system, where each cell expresses both Na_V_1.5 and K_ir_2.1 channels. The interfacial system has a spontaneously active phase (blue region) which is absent in the homogenous system. Orange star corresponds to the parameters used for the simulations in (B). D) Steady-state resting voltage and sodium channel availability along a line perpendicular to the interface. The voltage varies continuously between a polarized state on the K_ir_2.1 side and a depolarized state on the Na_V_1.5 side. The available reserve of Na_V_1.5 conductance (g_Na_ × (1 – m^3^) h) has a peak near the interface. The band of high Na_V_1.5 reserve supports AP propagation. Scale bar 0.2 mm. (E) Numerical simulation of simplified FHN-like model of topological action potentials (see Supplementary Model 2 for details). Left: Image of a topological action potential propagating along a tissue interface. Right: kymograph showing topological action potential propagation (Movie 4). See also Figs. S3-S5.

To better understand our results, we simulated a conductance-based model of our experiment using established parameters for K_ir_2.1 and Na_V_1.5^21,35^ (Supplementary Model 1). This model reproduced the basic phenomena, showing interface-localized action potential initiation and propagation (Fig. 2B, Movie 3). We then simulated the effect of changes in the expression level of the gap junctions, Na_V_1.5 or K_ir_2.1. Modulating the gap junction conductance, G_gj_, changed the width, length, and conduction speed of the topological action potential, with all three parameters scaling as (G_gj_)^1/2^, as one would expect from dimensional analysis (Fig. S3). Topological action potentials arose over at least a 60-fold variation in *g*_Na_ and at least a 1000-fold variation in *g*_K_, persisting to the highest levels of *g*_K_ and *g*_Na_ simulated (Fig. 2C).

We then performed analogous simulations where the Na_V_1.5 and K_ir_2.1 were homogeneously distributed through all cells. For cells co-expressing both ion channels, excitability occurred over a much narrower range of expression levels (∼30-fold in Na_V_1.5, but only ∼3-fold in K_ir_2.1; Fig. 2C). In the case of homogeneous expression, excitability required a delicate balance of depolarizing (Na_V_, leak, channelrhodopsin) and polarizing (K_ir_) currents to drive all the phases of the action potential. In contrast, in the interfacial structure, gap junction-mediated currents caused the relative contributions of Na_V_ and K_ir_ to vary from 0 to 100% transverse to the interface, dramatically expanding the range of parameters which supported excitability (Fig. 2D).

To facilitate an understanding of the basic requirements for topological action potentials, we developed a simplified model inspired by the FitzHugh-Nagumo (FHN) model (Supplementary Model 2; Supplementary Code).^36,37^ We broke the characteristic N-shaped nonlinearity of the FHN model into two components: one approximated the shape of a Na_V_ current-voltage (*I*-*V*) curve, and one approximated the shape of a K_ir_ *I*-*V* curve. These two components, each active in only one half-space, each had a single zero-crossing and hence a single stable resting potential. At the interface where both currents contributed, the *I-V* curve had three zero crossings and hence could support nonlinear excitations (Fig. S4). The simplified FHN-like model reproduced our observed propagating topological action potentials (Fig. 2E). Changes to the model parameters also yielded many other types of topological excitations, with diverse voltage patterns in space and time (Fig. S5C-E; Movie 4). These observations suggest that tissue interfaces might support a rich repertoire of novel bioelectrical phenomena.

We then explored experimentally the interaction of interface topology and sample topology. Using a plastic drinking straw as a separator, we plated Na_V_1.5-expressing cells in a disk and K_ir_2.1-expressing cells in the surrounding region (Fig. 3, Methods). We then removed the straw and let the cells form a circular interface. Optical stimulation in a line across the interface evoked topological action potentials which propagated bidirectionally around the interface, annihilating each other when they collided. To evoke a directional wave, we applied a tonic stimulation with a line to create a thin zone of Na_V_ channel inactivation. We then turned off the tonic stimulation and stimulated briefly to one side of this zone. This stimulus protocol created an action potential propagating in only one direction. The inactivated Na_V_ channels recovered by the time the action potential completed one trip around the disk. Thereafter the action potential propagated around the interface, circulating stably for many minutes (Fig S6; Movie 5). These results demonstrate that topological action potentials can sense the topology of tissue boundaries and can store information via their handedness of circulation.

**Figure 3.**
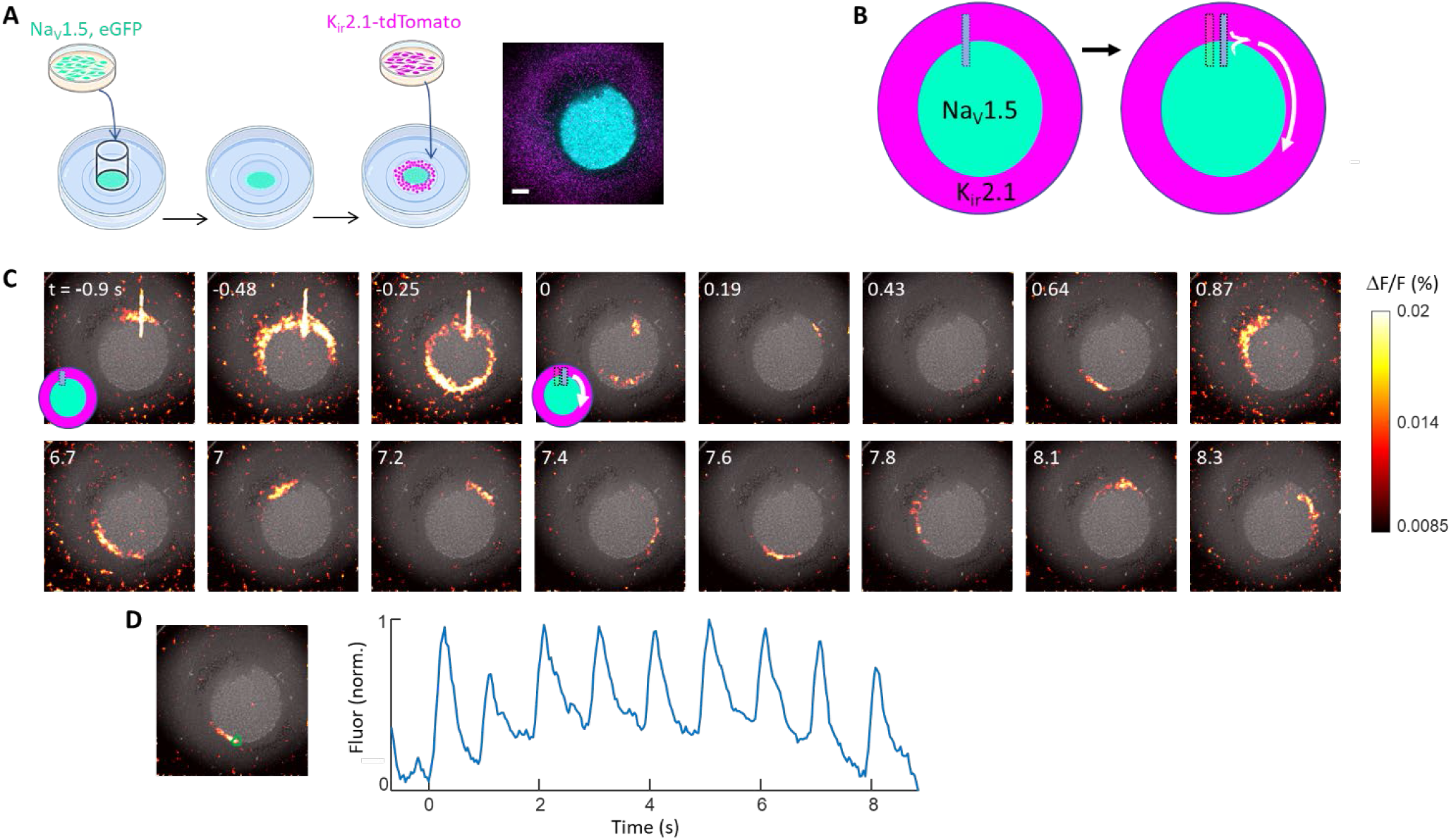
Persistently propagating topological action potentials at a circular interface. A) Sample preparation. HEK cells expressing Na_V_1.5 and CheRiff were seeded into circular region defined by a thin plastic cylinder. HEK cells expressing K_ir_2.1 and CheRiff were then added and settled in the unoccupied periphery. B) Stimulation protocol for evoking a unidirectional topological action potential wave. A long (450 ms) illumination pulse was delivered to a portion of the interface, to drive local Na_V_1.5 inactivation. A short (∼30 ms) pulse was then delivered to a nearby interface region, evoking a topological action potential that propagated unidirectionally. By the time the action potential completed one cycle, the inactivated Na_V_ channels had recovered, permitting cyclical propagation. C) Montage showing application of the inactivation stimulus, the trigger stimulus, and persistent AP propagation along the interface. D) Time-dependent fluorescence in the region indicated by the green polygon, showing a persistently propagating topological action potential. Scalebars: 1 mm. See also Fig. S6; Movie 5.

In the presence of voltage-gated Ca^2+^ channels, topological excitations might lead to locally elevated Ca^2+^, which could then turn on downstream patterns of gene expression. First, we tested whether co-expression of a T-type calcium channel, Ca_V_3.2, and K_ir_2.3 in HEK cells were sufficient, on their own, to support action potential propagation. We used HEK cells with constitutive expression of K_ir_2.3, doxycycline-inducible expression of Ca_V_3.2,^38^ and lentiviral expression of CheRiff for optogenetic stimulation. We induced Ca_V_3.2 expression with doxycycline and then mapped the voltage in confluent monolayer cultures of these cells (Fig. S7A). Local optogenetic stimulation induced action potentials which propagated radially outward and then broke up into self-sustaining spiral waves (Movie 6). Incubation with the Ca_V_ blocker nifedipine abolished the excitability (Fig. S7B). Samples where Ca_V_3.2 expression was not induced were also not excitable (Fig. S7C). Together, these results establish that Ca_V_3.2 and K_ir_ 2.3 are sufficient to sustain propagating action potentials in confluent HEK cell monolayers.

We then asked whether topological action potentials could arise at the interface between cells expressing Ca_V_3.2 and cells expressing K_ir_2.1. As with the Na_V_ channels, optogenetic stimulation with a stripe that spanned the interface induced excitations in membrane voltage which propagated along the interface (Fig. 4A, Movie 7). In separate samples, we used a far-red calcium-sensitive dye (Methods) and observed propagating Ca^2+^ signals along the interface too (Fig 4B, Movie 8). These results show that topological action potentials can be driven by multiple types of depolarizing channels (Na_V_, Ca_V_) and can drive increases in intracellular Ca^2+^ in cells within an electrical coupling length of a tissue interface.

**Figure 4.**
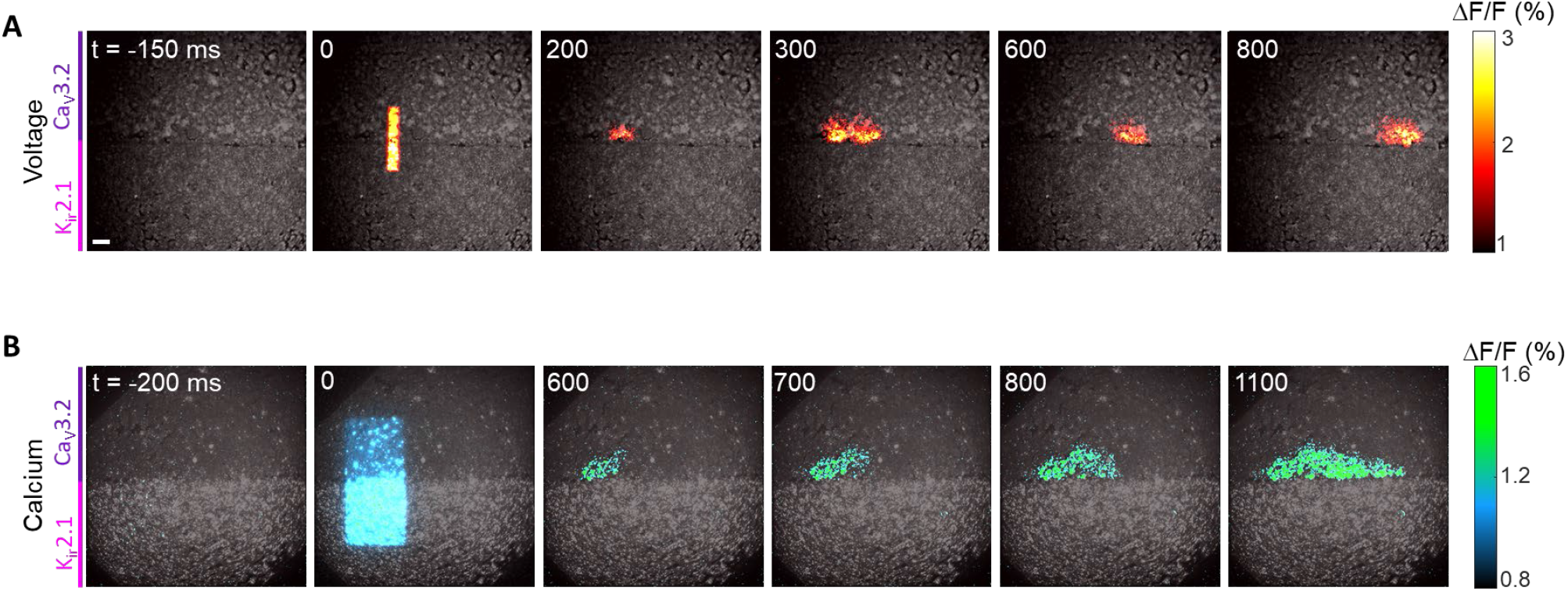
Topological action potentials driven by Ca^2+^ currents. A) Montage of frames from a voltage-imaging movie (Movie 7). Optogenetic stimulation spanning the interface at *t* = 0 evoked a topological action potential that propagated outward solely along the interface. B) Montage of frames from a calcium-imaging movie (from a different sample, Movie 8). Optogenetic stimulation at the interface evoked a calcium signal that propagated along the interface. Scale bar 1 mm. Baseline signal in gray, *ΔF*/*F* in color. See also Fig. S7.

## Discussion

The nonlinear behavior of ion channels, together with gap junctional sharing of currents, gives tissue interfaces unique excitability properties different from the bulk. Here we showed that mixtures of non-excitable cells can be excitable; an interface between non-excitable tissues can be excitable; a circular tissue interface can support stably circulating topological excitations; and topological excitations can drive local elevations in Ca^2+^ concentration.

The existence of topological action potentials at tissue interfaces raises the possibility of other types of topological effects in electrophysiology. For instance, our numerical simulations predicted a regime in which a Na_V_1.5-K_ir_2.1 interface showed spontaneous spiking, whereas cells co-expressing Na_V_1.5 and K_ir_2.1 did not show spontaneous spiking under any conditions. Other combinations of ion channels could lead to additional topological effects. For instance, expression of pacemaker channels on one side of the interface might lead to spontaneous spiking that starts only when two tissues touch, providing a means to signal first contact. Depending on the other ion channels on both sides of the interface, these contact-evoked spikes could either propagate just along the interface, into the bulk on one side, or into the bulk on both sides. Topological electrophysiology mechanisms could enable cells to sense the presence of nearby tissue interfaces, to determine the topology of their tissue environment, or to communicate over long distances.

The ways in which bioelectrical signaling complements chemical signaling pathways in embryonic development and morphogenesis^1,39^ are still being worked out. It is not yet known whether topological excitations arise in native tissues. Recent work has highlighted the importance of voltage-gated Ca^2+^ channels in development.^40–42^ These effects occur in nominally non-excitable cells, though our results highlight that excitability is not an intrinsic feature of a cell but rather depends on the surrounding tissues. Interactions between zebrafish skin melanophores and xanthophores also suggest a contact-dependent bioelectrical signaling mechanism.^43^ Our results suggest that analyses of such effects should not just examine the properties of individual cell types, but should also consider tissue interfaces as distinct electrophysiological entities. Finally, we note that due to the parallel structure of the Hodgkin-Huxley and Turing reaction diffusion equations, analogous interfacial excitations could also occur in chemical reaction-diffusion systems. More broadly, topological action potential could serve as a testbed for investigating emergent phenomena at interfaces of nonlinear interacting elements.

## Supporting information

Matlab code for FitzHugh-Nagumo-type simulations

Movie 1: Propagating action potential in a mixed monolayer of cells expressing either NaV1.5 or Kir2.1.

Movie 2: Topological action potential at a linear interface.

Movie 3: Conductance-based simulation of a topological action potential.

Movie 4: Diverse topological excitations in a FHN-like model.

Movie 5: Topological action potential at a circular interface.

Movie 6: Propagating action potential in cells co-expressing CaV3.2, Kir2.3, and CheRiff.

Movie 7: Topological action potential at a Kir2.1|CaV3.2 interface (Voltage signal).

Movie 8: Topological action potential at a Kir2.1|CaV3.2 interface (Ca2+ signal).

## Acknowledgments

This work was supported by the Vannevar Bush Faculty Fellowship (A.E.C.), a National Science Foundation Quantum Leap Challenge Institute on Quantum sensing for Biophysics and Bioengineering (QuBBE) (A.E.C.) and an EMBO Fellowship to H.O (ALTF 543-2020). We thank Noam Ziv and his laboratory for hosting H.O. during the COVID-19 pandemic. We thank Terrance Snutch for helpful discussions and for providing cells expressing Ca_V_3.2 and K_ir_2.3. We thank Evan Miller for the BeRST1 dye. We thank Vincenzo Vitelli, Colin Scheibner and Suyang Xu for helpful discussions. We thank Shahinoor Begum, Andrew Preecha, Tamar Galateanu and Lonya Odessky for technical assistance.

## Methods

### Cell line generation and culture

Cells stably co-expressing Na_V_1.5 and CheRiff-eGFP were described in Ref ^31^. For cells expressing K_ir_2.1 and CheRiff, cells constitutively expressing CheRiff were transiently transfected with pCAG-Kir2.1-T2A-tdTomato (from Massimo Scanziani, Addgene 60598) using CalFectin transfection reagent (SignaGen Laboratories) following manufacturer’s directions. Cells were seeded in imaging plates 24-48 hours after transfection. Cell culture maintenance was the same as for Optopatch Spiking HEK cells^34^. Cells stably expressing human Ca_V_3.2, along with cells constitutively expressing K_ir_2.3 and doxycycline inducible Ca_V_3.2 were a generous gift from Terrance Snutch and are described in Ref. ^38^. These cells were infected with a lentiviral vector expressing pCMV-CheRiff-CFP (Addgene 136636) to render them light sensitive.

### Sample preparation

35 mm dishes with 14 mm glass coverslip bottoms (Cellvis, #D35-14-1.5-N) were coated with polyethyleneimine (PEI). Two days prior to imaging, linear interfaces were created by seeding Na_V_1.5- and K_ir_2.1-expressing cells separated by a 100 µm thick plastic divider, 4×10^5^ cells of each type. After 2 h, the divider was carefully removed, the plate was gently washed with phosphate-buffered saline (PBS) to remove unattached cells, the plate was incubated in culture medium. For the circular interfaces, the central island was created by seeding 5×10^5^ Na_V_1.5-expressing cells inside a region defined by a circular 5 mm diameter plastic cylinder (a cut drinking straw). After 2 h, the cylinder was removed, unattached cells were removed by gentle washing with PBS, and the plate was incubated overnight with culture medium. Approximately 10^6^ K_ir_2.1-expressing cells were then seeded onto the dish. The K_ir_2.1 cells primarily attached in the annular region not already occupied by Na_V_1.5-expressing cells.

### Imaging and stimulation

For voltage imaging, cells were washed to remove culture medium and then incubated with 1-2 μM BeRST1 dye in PBS for 30 min in a tissue culture incubator. Immediately before imaging, samples were washed twice and immersed in imaging solution containing (in mM) 125 NaCl, 15 HEPES, 25 glucose, 2.5 KCl, 1 MgCl_2_, 2 CaCl_2_ with pH adjusted to 7.3 with NaOH. Sample preparation for Ca^2+^ imaging was similar, except that the cells were washed and incubated with 3 μM BioTracker™ 609 Red Ca^2+^ AM Dye (EMD Millipore #SCT021) in PBS for 30 min in a tissue culture incubator, then washed and immersed in imaging solution. Widefield imaging was performed using an Axiovert 200 (Zeiss) inverted microscope, equipped with a light source (xenon arc, XBO 75 W) and filter sets for the GFP, tdTomato, BeRST1 and calcium dye channels. Light was collected using a 2X, numerical aperture 0.1 objective lens (ThorLabs #TL2X-SAP) into an EMCCD camera (Andor iXon EM+ 897, 512×512 pixels, 35 frames/s). The field of view (FOV) was expanded to 9.9 mm in the sample plane by placing a 0.5X image reducer (View Solutions, #MA513302) in the light collection path. CheRiff was excited by a 470 nm LED (ThorLabs #M470L4-C4) connected to the topside illumination path. The stimulation region was defined by a thin slit inserted in the excitation beam and projected onto the sample plane via the microscope condenser. The camera was controlled via the Andor SOLIS software. The LED illumination was controlled via a National Instruments DAQ card. Camera and LED were synchronized using a custom Matlab code. For imaging and patterned optogenetic stimulation of Ca_V_3.2-K_ir_2.3 monolayers and Ca_V_3.2-K_ir_2.1 interfaces we used an ultrawidefield microscope as, described in Ref. ^44^.

### Data analysis

All data were processed and analyzed using custom software (MATLAB). The baseline fluorescence, *F*, of the BeRST1 signal was calculated for each pixel by the mean intensity over the time series. The movie was then converted into units of ΔF/F and smoothed by spatially filtering with a 12×12 pixels kernel. For the linear interface experiments (Fig. 1) 10 consecutive trials were averaged before converting to ΔF/F units. To calculate the spatial shape of the propagating AP (Fig. 2A), images acquired at different times were laterally shifted to cancel the motion of the AP wavefront. Background noise from variation between rows was suppressed by setting pixels below the 20^th^ percentile of their row to the minimum intensity of the frame. Finally, the movie was averaged along time with a 12 frames window.

Kymographs were constructed by averaging rectangles of 40×10 pixels along the interface. For the circular interface experiment (Fig. 3) the time series of the ΔF/F movie was reconstructed from a detrended vectorized ΔF/F sequence to account for changes in baseline signal due to photobleaching. In addition to spatial filtering movies were averaged along time with a 6 frame sliding window. For the time trace plot (Fig. 3D) a spatial region of interest was defined and averaged across pixels.

GFP and tdTomato images (Fig. 1A,B) were processed by subtracting a background image (no illumination) and dividing pixel-wise by a reference image (dish with no cells) to account for spatial variations in illumination intensity. A gamma correction was applied to the tdTomato image to permit visualization of cells with widely varying expression levels.

### Numerical modeling

Model equations and simulation methods can be found in Supplementary Models 1 and 2, and Supplementary Code.

## Supplementary Material

This file contains:

Supplementary models 1 and 2

Supplementary figures S1-S7

Supplementary movie captions 1-8

### Supplementary Model 1: Hodgkin-Huxley-like model of interface APs

For realistic simulations of tissue interfaces, we used a Hodgkin-Huxley (HH)-like model, with the Na_V_1.5 and K_ir_2.1 currents separated along the x axis:

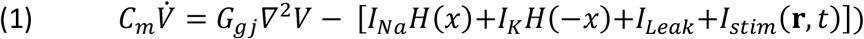

where *H* (*x*) is the Heaviside step function:

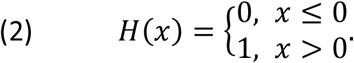

Details of the simulation approach follows previous work from our lab.^34^ *I*_*Na*_ represents a Na_V_1.5 current, modeled using the Ten-Tusscher model.^45^ The slow inactivation gate was set to *j* = 1, and the fast *m* gate was replaced by its asymptotic value *m*_*∞*_(*V*):

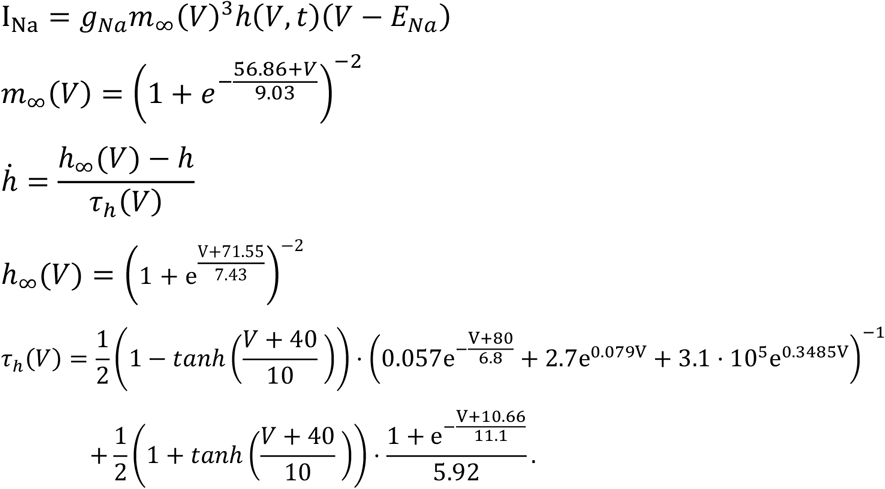

Here τ_*h*_(*V*) is a smoothed version of the original function in the Ten-Tusscher model, for numerical stability. The sodium reversal potential was set to *E*_*Na*_ = 75 mV.

*I*_*K*_ represents current through the inward-rectifying K_ir_2.1 channel, modeled as an instantaneous function of voltage using the Ten-Tusscher model:^45^

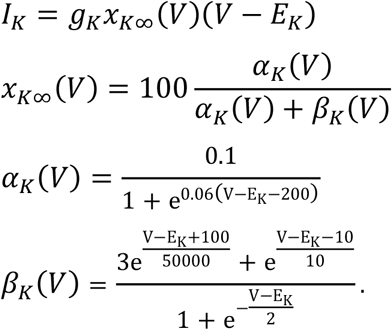

The potassium reversal potential was set to *E*_*K*_ = -107 mV.

The leakage term is an Ohmic conductance with resting potential at *V* = 0:

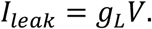

*I*_*stim*_(***r, t***) is a stimulation term, *g*_*ChR*_ (**r, *t***)*V*, patterned in space and time to mimic optogenetic stimulation.

All conductances were expressed in units of 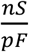, so *C*_*m*_ = 1. Spatial units were set to the spatial dimensions of a single cell, hence the gap-junction conductance *G*_*gj*_ had the same units as the ionic conductances. All voltages were expressed in mV and time in ms.

Simulations were performed in Matlab with a 300×300 grid of cells assigned to express either *I*_*Na*_or *I*_*K*_, depending on each cell’s location. Cell size and capacitance were set to 10 µm and 10 pF respectively. Gap-junction conductance was set to G_gj_ = 10 nS/pF. An Ohmic leak conductance, g_L_ = 0.005 nS/pF, *E*_*Leak*_ = 0, was added to maintain stable resting potential in all cells. CheRiff stimulation was modeled as an additional Ohmic conductance with *E*_*ChR*_ = 0 and g_ChR_ = 0.5 nS/pF during periods of illumination. To calculate gap junctional currents, the Laplacian of the voltage was approximated by the discrete second-difference. No-flux boundary conditions were applied at the edges of the simulation. Euler’s method was implemented for time integration with dt = 0.01 ms.

### Supplementary Model 2: FitzHugh-Nagumo (FHN)-inspired model of interface APs

The purpose of the model presented hereafter in Equations (5) and (6) is to capture the key physics of excitable tissue interfaces in simplified equations which are more analytically tractable than the Hodgkin-Huxley (HH)-like model presented above.

As an intermediate step, we simplify the HH-like model while maintaining the original parameters by replacing the complicated functions *x*_*K∞*_(*V*) and *h*_*∞*_(*V*) by simple exponential functions of *V* with shapes that are qualitatively similar within the relevant voltage range, and replacing τ_*h*_(*V*) by a constant:

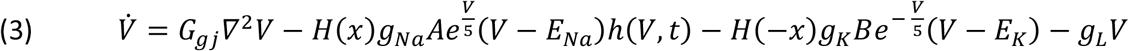

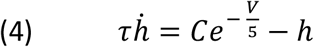

where A, B, C, and *τ* are constants, and *H*(*x*) is the Heaviside function (Equation (2)). This model reproduced the topological action potentials of the HH-like model with the values: 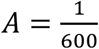, *B* = 3 · 10^−7^, *C* = 3 · 10^−8^, *τ* = 50.

To further simplify the model, we adapted the FitzHugh-Nagumo (FHN) model^36,37^ of neuronal excitability. We rescaled the voltage variable of Eq. (3), introduced a linear equation for the inactivation variable *h*, and dropped the leakage term. The fully simplified model is:

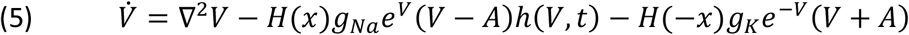

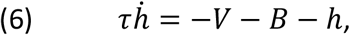

where *H*(*x*) is the Heaviside function (Equation (2)). Here we express the conductances *g*_*Na*_ and *g*_*K*_as unitless entities, representing ratios to the gap-junction conductance, which we set to 1.

In contrast to the FHN model, in our model the current term 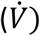 retains the overall structure of the HH model,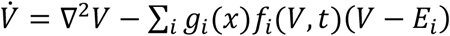. Therefore, its parameters retain, at least approximately, a biophysical interpretation. In Eq. (5), the second term on the r.h.s. corresponds to a Na_V_-like current, and the third term corresponds to a K_ir_-like current, though these terms no longer match specific ion channels.

In the presence of both K_ir_ and Na_V_ currents (i.e. near the interface), the I-V curve (or 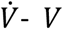 curve) is designed to be ‘N’ shaped, similar to the FHN model. But for K_ir_-only or Na_V_-only cells (i.e. far from the interface), the I-V curve only has a single zero-crossing (Fig. S4). This arrangement promotes excitability on the interface only, where gap junctional coupling mediates sharing of currents across the interface. The ‘N’ shaped curve fulfills the conditions for excitability, as it has 2 stable fixed points at *h* = 1. Starting at the lower (more polarized) fixed point, a depolarizing stimulus engages positive feedback which drives the system to the upper fixed point. The *h* variable then starts to decrease, mimicking Na_V_ channel inactivation. At a critical lower value of the *h* variable, the upper fixed point disappears via a saddle-node bifurcation, forcing the system to repolarize and return to the lower fixed point.

This simple model produced a rich repertoire of interface excitability phenomena. Qualitatively distinct dynamics can be produced by changing the model parameters. The closest approximation to the experimental results (Fig. 2E) was obtained with the following parameters: *g*_*Na*_= 10^−2^,*g*_*K*_ = 10^−5^, *τ* = 10^4^, *A* = 5, *B* = 3. In Fig. S5, we consider the parameters *g*_*Na*_= 10^−2^; *τ* = 10^4^; *A* = 5; *B* = 2, and vary the parameter *g*_*K*_. To test for excitability, we delivered the equivalent of an optogenetic stimulus, transiently introducing a current *g*_*ChR*_*V* along the left edge. As *g*_*K*_ increased, the system transitioned from non-excitable (Fig. S5A) to production of a single propagating AP (Fig. S5B), mimicking our experimental results. Further increases in *g*_*K*_led to a regime in which a single stimulus pulse switched the system from a quiescent state into a state where periodic impulses emerged from the stimulated zone, even after the stimulus was removed (triggered pacemaking, Fig. S5C). Other values of the parameters led to more exotic behaviors along the interface, such as coexistence of long-wavelength oscillations and impulse-like traveling waves (Fig. S5D), or emergence of slow timescales and irregularity (Fig. S5E, Movie 4). These kinds of dynamics did not occur in the spatially homogeneous versions of the model (where *g*_*k*_ and *g*_*Na*_ were spatially uniform), emphasizing the qualitatively distinct nature of interfacial excitability.

Matlab code for simulating the model is provided as Supplementary Code. Simulations were performed with time-steps of dt = 0.1 time units. Prior to each simulation the system was allowed to settle with the chosen set of parameters for 3000 time units, with the dynamics of the *h* gate quenched to those of its asymptotic value *h* = −*V* − *BB*. The full simulation was then executed, with a single stimulus delivered after an additional 3000 time units.

## Supplementary Figures

**Figure S1.**
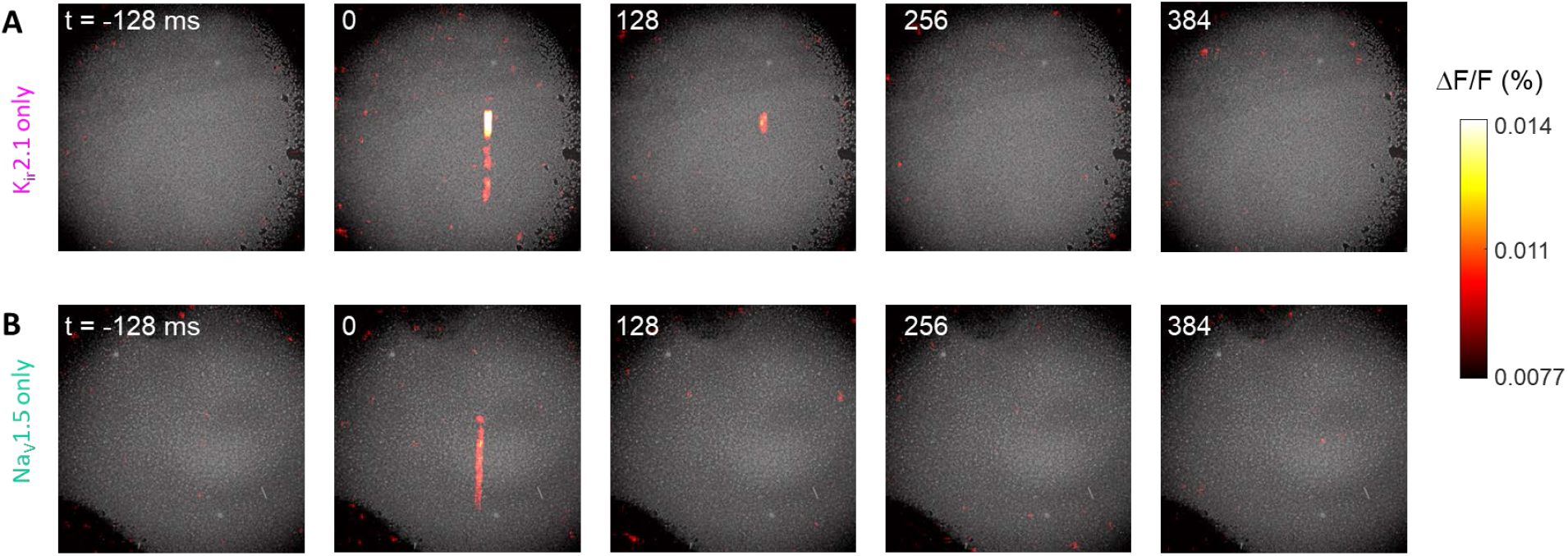
Control experiments showing absence of excitability in cells expressing only one voltage-dependent channel. In both panels, the cells also expressed a channelrhodopsin, CheRiff. The stimulus was delivered as a bar of blue light at *t* = 0. Related to Fig. 1.

**Figure S2.**
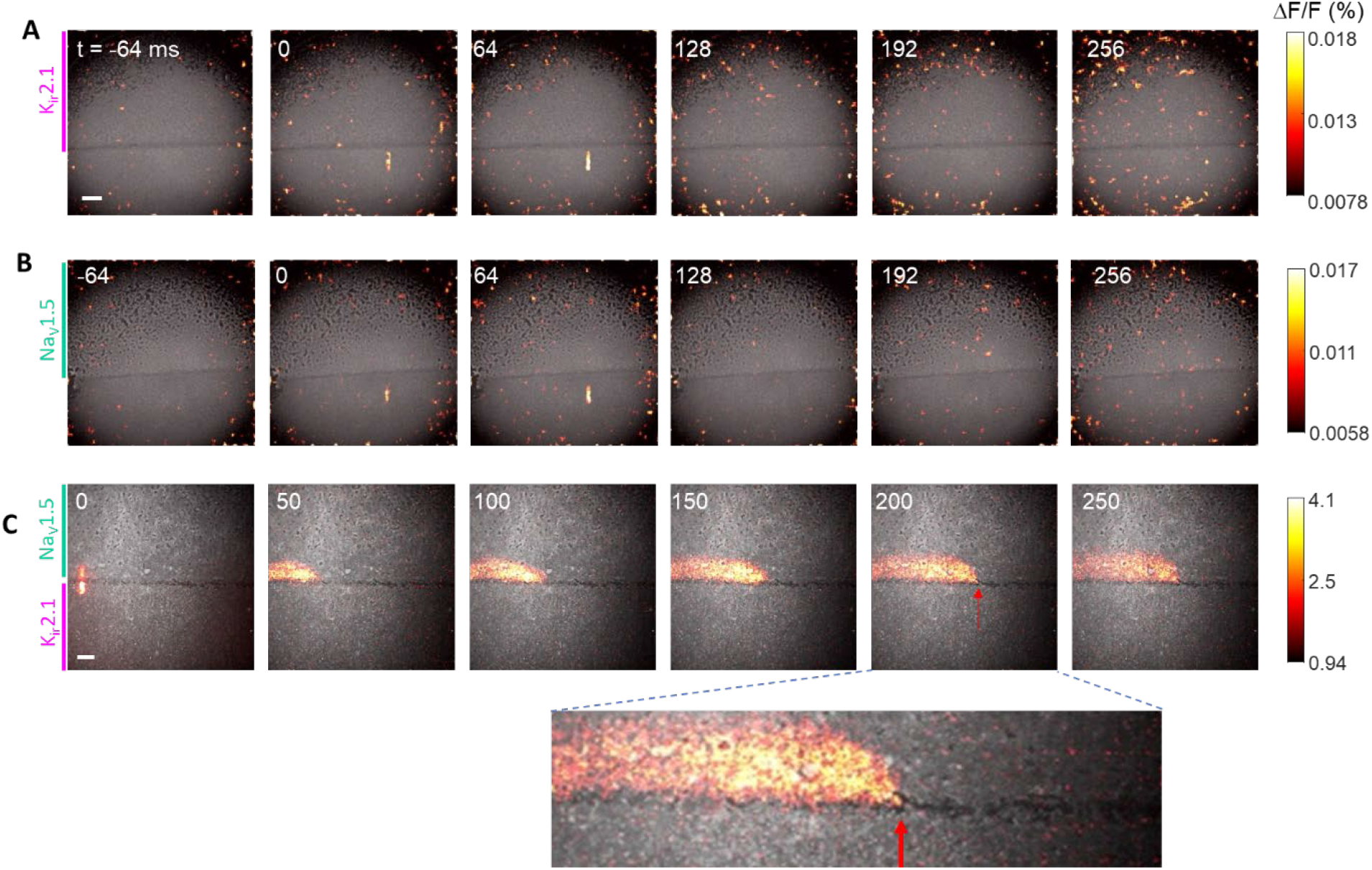
Control experiments establishing necessary conditions for topological action potentials. A,B) Interfaces between populations of K_ir_2.1- or Na_V_1.5-expressing cells and cells not expressing either ion channel are not excitable. C) Topological AP propagation in a Na_V_-K_ir_ interface was blocked in a region where cells did not migrate to fill the gap between the populations (red arrow). Scale bars 1 mm. Related to Fig. 1.

**Figure S3.**
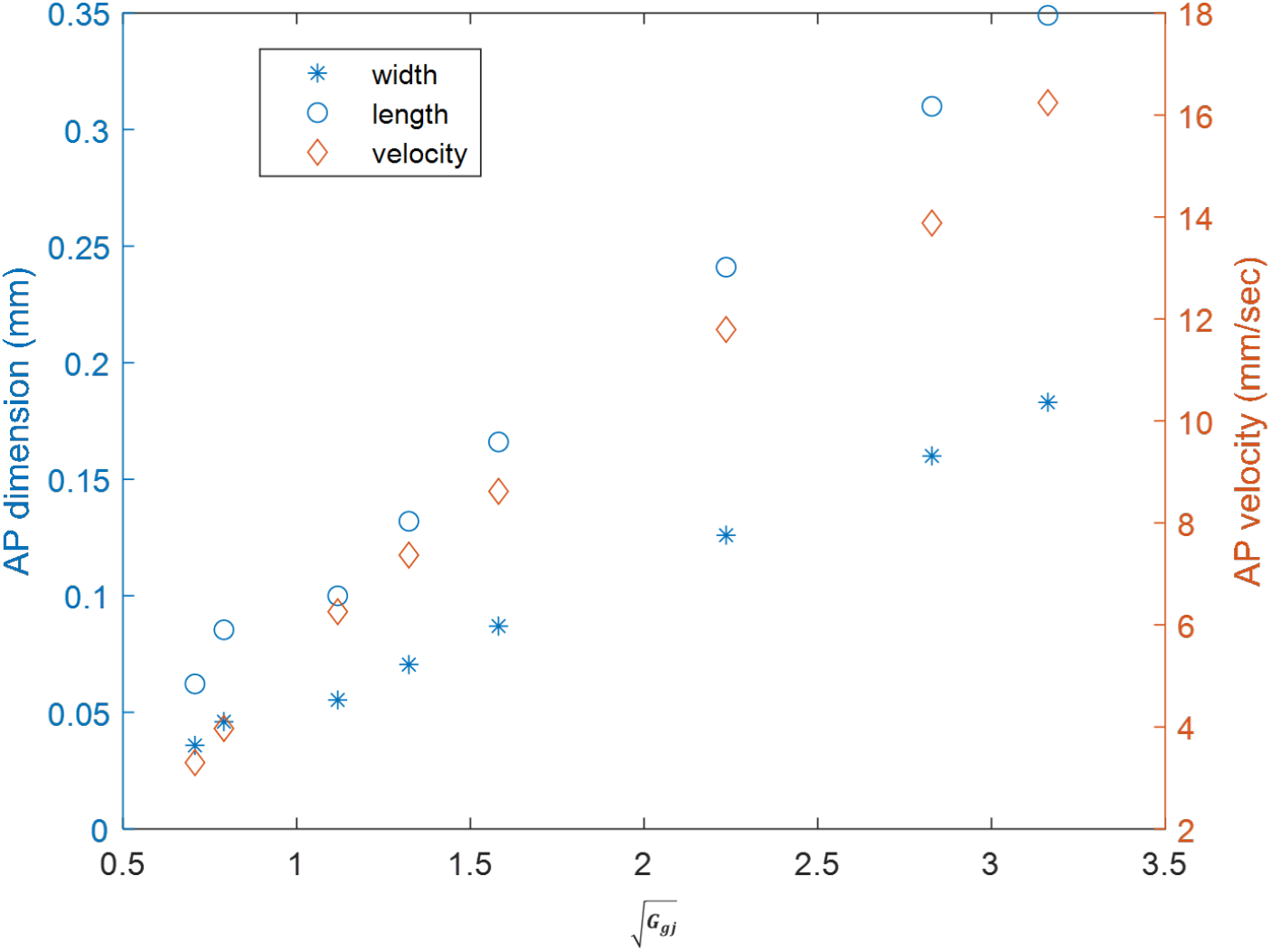
Effect of gap junction conductance on topological action potential dimensions and velocity. The left y-axis corresponds to the simulated width and length of the topological AP, measured at half peak. The right y-axis corresponds to the AP velocity, extracted from AP kymographs. As expected from dimensional analysis, all three quantities scale with 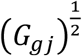. Related to Fig. 2.

**Figure S4.**
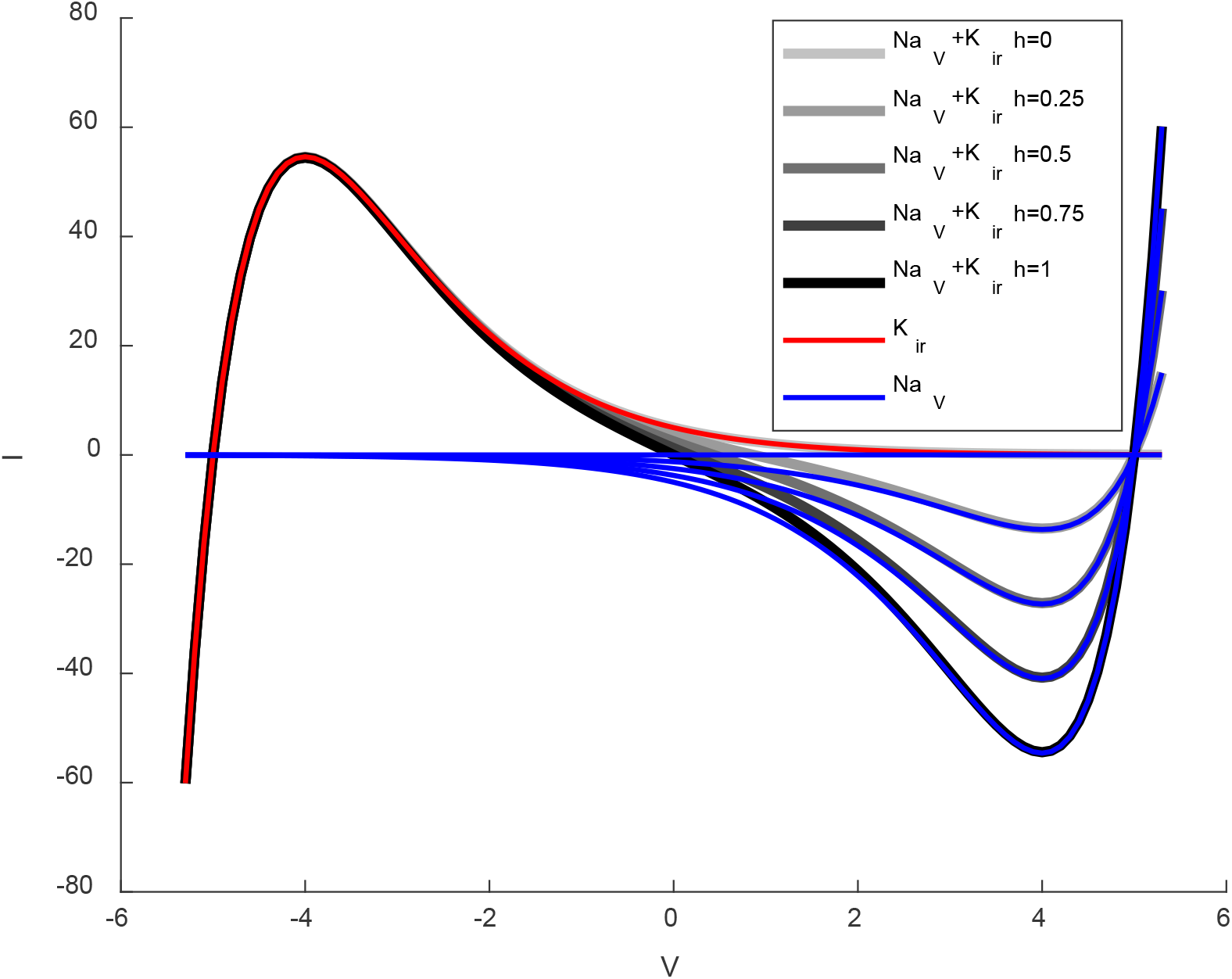
I-V curves of a FHN-like model. I-V curves of cells with equal amounts of K_ir_ and Na_V_ and different values of h (greyscale curves); a “K_ir_-only” cell (red); and “Na_V_-only” cells (blue). In this example, *A* = 5, g_Na_ = 1, g_K_ = 1, and gap junctional currents are omitted. The upper fixed point disappears via a saddle-node bifurcation at 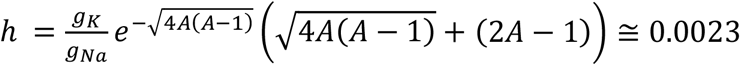. Related to Fig. 2.

**Figure S5.**
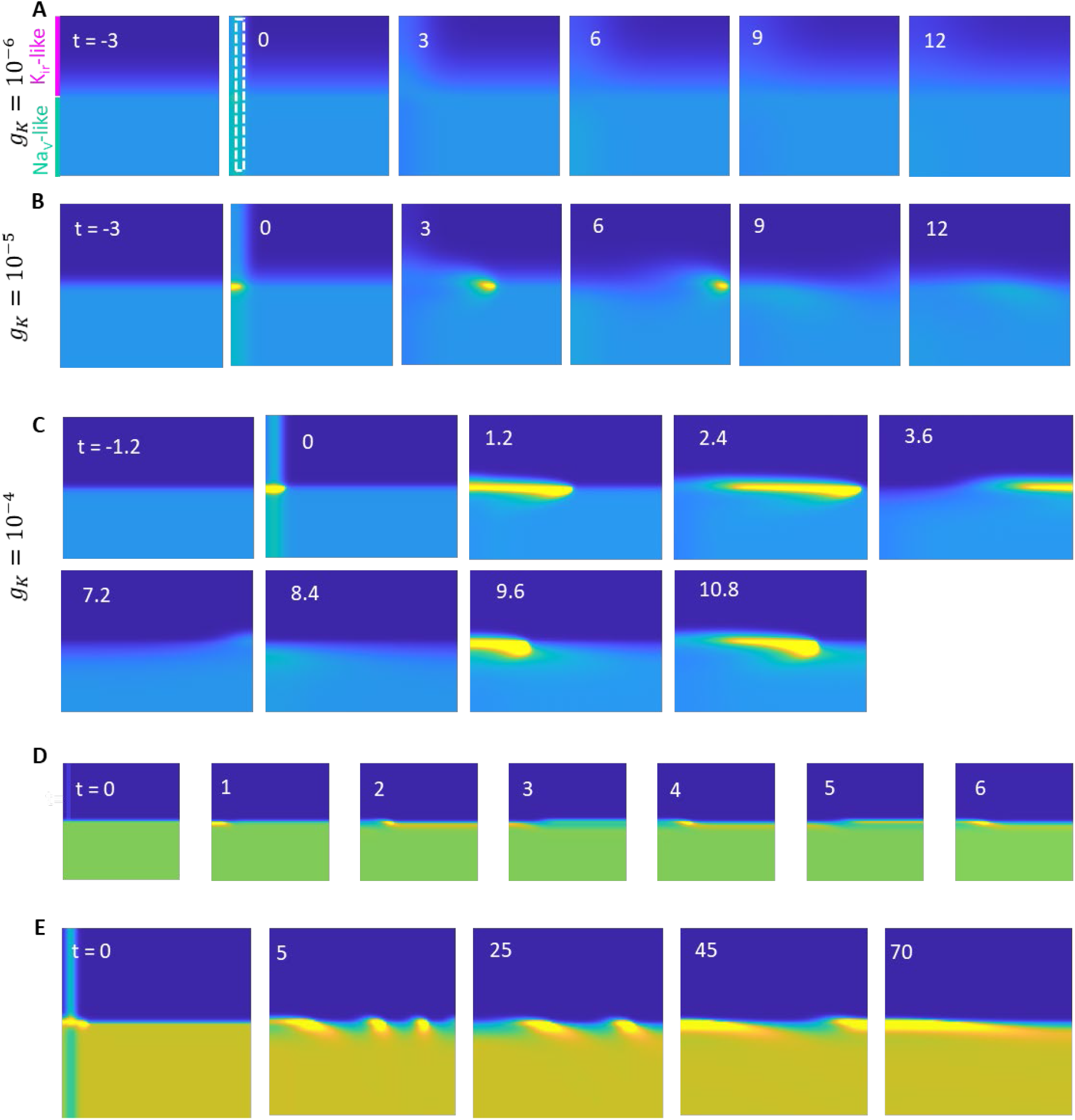
Interface AP propagation in the FitzHugh-Nagumo-inspired model. Increases in g_K_ led to transitions between A) non-excitability, B) single evoked AP, and C) triggered pacemaking. Other parameters were: *g*_*Na*_= 10^−2^; τ = 10^4^; *A* = 5; *B* = 2. Stimulus was delivered to a region 10 cells wide (marked by dotted line at t = 0) for 100 time units. D-E) Exotic interface dynamics. Parameters for D: *g*_*Na*_= 10^−1^; g_K_= 10^−3^; τ = 10^3^; *A* = 4; *BB* = 0. E) Same parameters except for *BB* = 1. Time stamps in all panels are in units of 1000 time points. Related to Fig. 2.

**Figure S6.**
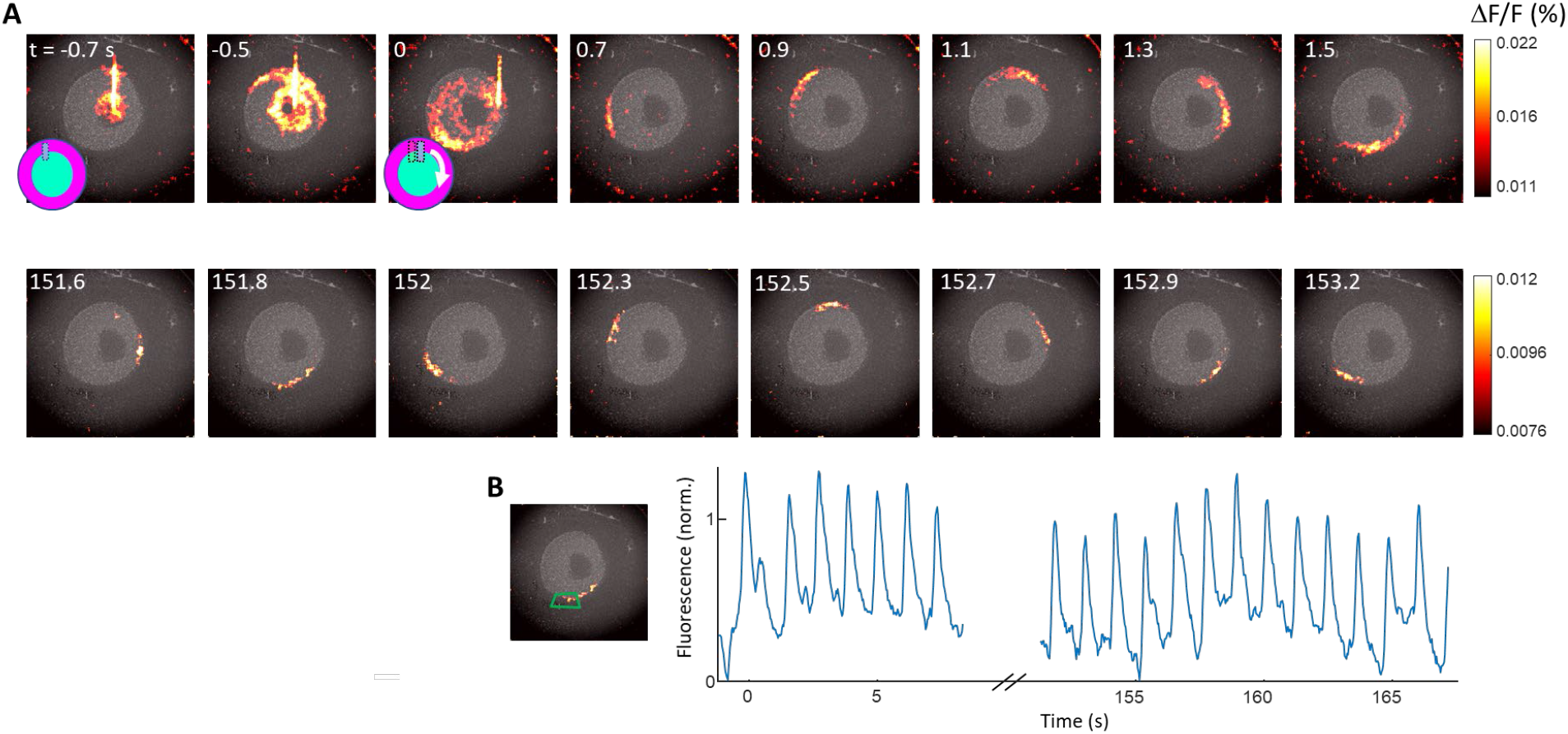
Topological action potentials can propagate along a circular interface. A) Montage showing application of the inactivation stimulus, the trigger stimulus, and AP propagating for > 150 s along the interface. B) Time-dependent fluorescence in the region indicated by the green polygon. Related to Fig. 3.

**Figure S7.**
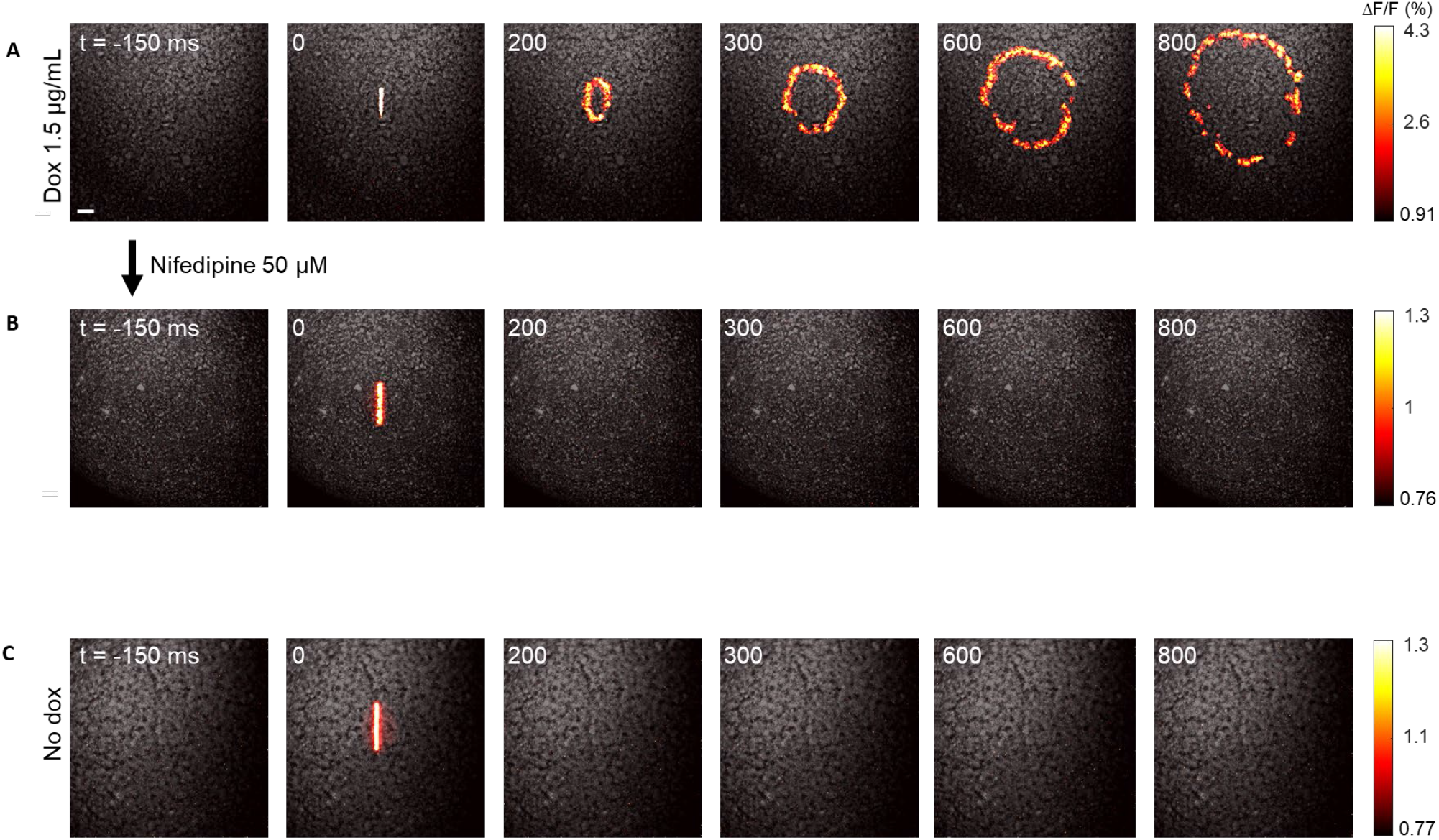
Calcium-driven action potentials. Monolayers of HEK cells were grown expressing Ca_V_3.2 (dox-induced), K_ir_2.3 and CheRiff. A) After dox application (1.5 *µ*g/mL for 1 day) to turn on Ca_V_3.2 channel expression, the monolayer supported optically evoked action potential wave propagation. Scale bar: 1 mm. B) The Ca_V_3.2 channel blocker nifedipine eliminated excitability of the cell monolayer. C) In the absence of doxycycline, the Ca_V_3.2 channel was not expressed and the monolayer was not excitable. Related to Fig. 4.

### Supplementary Movies

Movie 1: Propagating action potential in a mixed monolayer of cells expressing either Na_V_1.5 or K_ir_2.1.

Movie 2: Topological action potential at a linear interface.

Movie 3: Conductance-based simulation of a topological action potential.

Movie 4: Diverse topological excitations in a FHN-like model.

Movie 5: Topological action potential at a circular interface.

Movie 6: Propagating action potential in cells co-expressing Ca_V_3.2, K_ir_2.3, and CheRiff.

Movie 7: Topological action potential at a K_ir_2.1|Ca_V_3.2 interface (Voltage signal).

Movie 8: Topological action potential at a K_ir_2.1|Ca_V_3.2 interface (Ca^2+^ signal).

## References

1. Levin, M. Bioelectric signaling: Reprogrammable circuits underlying embryogenesis, regeneration, and cancer. Cell 184, 1971–1989 (2021).

2. Pietak, A. & Levin, M. Bioelectric gene and reaction networks: computational modelling of genetic, biochemical and bioelectrical dynamics in pattern regulation. J. R. Soc. Interface 14, 20170425 (2017).

3. Robinson, K. R. & Messerli, M. A. Left/right, up/down: the role of endogenous electrical fields as directional signals in development, repair and invasion. 8.

4. Levin, M. Endogenous bioelectrical networks store non-genetic patterning information during development and regeneration. J. Physiol. 592, 2295–2305 (2014).

5. Levin, M. Bioelectric mechanisms in regeneration: unique aspects and future perspectives. Semin. Cell Dev. Biol. 20, 543–556 (2009).

6. Tyler, S. E. B. Nature’s Electric Potential: A Systematic Review of the Role of Bioelectricity in Wound Healing and Regenerative Processes in Animals, Humans, and Plants. Front. Physiol. 8, 627 (2017).

7. Cervera, J., Alcaraz, A. & Mafe, S. Bioelectrical Signals and Ion Channels in the Modeling of Multicellular Patterns and Cancer Biophysics. Sci. Rep. 6, 20403 (2016).

8. Payne, S. L., Levin, M. & Oudin, M. J. Bioelectric Control of Metastasis in Solid Tumors. Bioelectricity 1, 114–130 (2019).

9. Hasan, M. Z. & Kane, C. L. Colloquium: Topological insulators. Rev. Mod. Phys. 82, 3045– 3067 (2010).

10. Vergniory, M. G. et al. A complete catalogue of high-quality topological materials. Nature 566, 480–485 (2019).

11. Haldane, F. D. M. & Raghu, S. Possible Realization of Directional Optical Waveguides in Photonic Crystals with Broken Time-Reversal Symmetry. Phys. Rev. Lett. 100, 013904 (2008).

12. Süsstrunk, R. & Huber, S. D. Observation of phononic helical edge states in a mechanical topological insulator. Science 349, 47–50 (2015).

13. Nash, L. M. et al. Topological mechanics of gyroscopic metamaterials. Proc. Natl. Acad. Sci. 112, 14495–14500 (2015).

14. Paulose, J., Chen, B. G. & Vitelli, V. Topological modes bound to dislocations in mechanical metamaterials. Nat. Phys. 11, 153–156 (2015).

15. Souslov, A., Dasbiswas, K., Fruchart, M., Vaikuntanathan, S. & Vitelli, V. Topological Waves in Fluids with Odd Viscosity. Phys. Rev. Lett. 122, 128001 (2019).

16. Gong, Z. et al. Topological Phases of Non-Hermitian Systems. Phys. Rev. X 8, 031079 (2018).

17. Dasbiswas, K., Mandadapu, K. K. & Vaikuntanathan, S. Topological localization in out-of-equilibrium dissipative systems. Proc. Natl. Acad. Sci. 115, E9031–E9040 (2018).

18. Murugan, A. & Vaikuntanathan, S. Topologically protected modes in non-equilibrium stochastic systems. Nat. Commun. 8, 13881 (2017).

19. Hou, J. H., Kralj, J. M., Douglass, A. D., Engert, F. & Cohen, A. E. Simultaneous mapping of membrane voltage and calcium in zebrafish heart in vivo reveals chamber-specific developmental transitions in ionic currents. Front. Physiol. 5, 344 (2014).

20. Adam, Y. et al. Voltage imaging and optogenetics reveal behaviour-dependent changes in hippocampal dynamics. Nature 569, 413 (2019).

21. McNamara, H. M. et al. Bioelectrical domain walls in homogeneous tissues. Nat. Phys. 16, 357–364 (2020).

22. Ma, Y. et al. Synthetic mammalian signaling circuits for robust cell population control. Cell 0, (2022).

23. Warmflash, A., Sorre, B., Etoc, F., Siggia, E. D. & Brivanlou, A. H. A method to recapitulate early embryonic spatial patterning in human embryonic stem cells. Nat. Methods 11, 847–854 (2014).

24. Cervera, J., Levin, M. & Mafe, S. Bioelectrical Coupling of Single-Cell States in Multicellular Systems. J. Phys. Chem. Lett. 11, 3234–3241 (2020).

25. Xu, J. et al. The Role of Cellular Coupling in the Spontaneous Generation of Electrical Activity in Uterine Tissue. PLOS ONE 10, e0118443 (2015).

26. Agarwala, A. & Shenoy, V. B. Topological Insulators in Amorphous Systems. Phys. Rev. Lett. 118, 236402 (2017).

27. Pöyhönen, K., Sahlberg, I., Westström, A. & Ojanen, T. Amorphous topological superconductivity in a Shiba glass. Nat. Commun. 9, 2103 (2018).

28. Mitchell, N. P., Nash, L. M., Hexner, D., Turner, A. & Irvine, W. T. M. Amorphous topological insulators constructed from random point sets. Nat. Phys. 14, 380–385 (2018).

29. Park, J. et al. Screening fluorescent voltage indicators with spontaneously spiking HEK cells. PloS One 8, e85221 (2013).

30. McNamara, H. M., Zhang, H., Werley, C. A. & Cohen, A. E. Optically controlled oscillators in an engineered bioelectric tissue. Phys. Rev. X 6, 031001 (2016).

31. Zhang, H., Reichert, E. & Cohen, A. E. Optical electrophysiology for probing function and pharmacology of voltage-gated ion channels. eLife 5, e15202 (2016).

32. Hochbaum, D. R. et al. All-optical electrophysiology in mammalian neurons using engineered microbial rhodopsins. Nat. Methods 11, 825–833 (2014).

33. Huang, Y. L., Walker, A. S. & Miller, E. W. A photostable silicon rhodamine platform for optical voltage sensing. J. Am. Chem. Soc. 137, 10767–10776 (2015).

34. McNamara, H. M. et al. Geometry-dependent arrhythmias in electrically excitable tissues. Cell Syst. 7, 359-370. e6 (2018).

35. ten Tusscher, K.H.W.J., Noble, D., Noble, P. J. & Panfilov, A.V. A model for human ventricular tissue. Am. J. Physiol.-Heart Circ. Physiol. 286, H1573–H1589 (2004).

36. FitzHugh, R. Impulses and physiological states in theoretical models of nerve membrane. Biophys. J. 1, 445–466 (1961).

37. Nagumo, J., Arimoto, S. & Yoshizawa, S. An active pulse transmission line simulating nerve axon. Proc. IRE 50, 2061–2070 (1962).

38. Belardetti, F. et al. A Fluorescence-Based High-Throughput Screening Assay for the Identification of T-Type Calcium Channel Blockers. ASSAY Drug Dev. Technol. 7, 266–280 (2009).

39. Gregor, T., Tank, D. W., Wieschaus, E. F. & Bialek, W. Probing the Limits to Positional Information. Cell 130, 153–164 (2007).

40. Pitt, G. S., Matsui, M. & Cao, C. Voltage-Gated Calcium Channels in Nonexcitable Tissues. Annu. Rev. Physiol. 83, 183–203 (2021).

41. Atsuta, Y., Tomizawa, R. R., Levin, M. & Tabin, C. J. L-type voltage-gated Ca2+ channel CaV1.2 regulates chondrogenesis during limb development. Proc. Natl. Acad. Sci. 116, 21592– 21601 (2019).

42. Lin, S.-S. et al. Cav3.2 T-type calcium channel is required for the NFAT-dependent Sox9 expression in tracheal cartilage. Proc. Natl. Acad. Sci. 111, E1990–E1998 (2014).

43. Inaba, M., Yamanaka, H. & Kondo, S. Pigment pattern formation by contact-dependent depolarization. Science 335, 677 (2012).

44. Werley, C. A., Chien, M.-P. & Cohen, A. E. Ultrawidefield microscope for high-speed fluorescence imaging and targeted optogenetic stimulation. Biomed. Opt. Express 8, 5794–5813 (2017).

45. en Tusscher, K. H. W. J., Noble, D., Noble, P. J. & Panfilov, A. V. A model for human ventricular tissue. Am. J. Physiol. -Heart Circ. Physiol. 286, H1573–H1589 (2004).

